# Quantification of small molecule partitioning in a biomolecular condensate with 2D NMR spectroscopy

**DOI:** 10.1101/2025.05.25.656010

**Authors:** Julie Maibøll Buhl, Sujata Mahapatra, Magnus Kjærgaard, Frans A. A. Mulder

## Abstract

Several intrinsically disordered proteins have been shown to undergo phase separation into a dense and dilute phase and this process is intimately linked with the regulation of cellular processes. It is therefore highly relevant to know how metabolites partition between these phases. We show here that the partitioning of components in a complex mixture can be robustly obtained from a single set of 2D NMR spectra recorded on the dilute and dense phases separately using a so-called “time zero extrapolated approach” known as HSQC_0_. The spectral separation power of 2D NMR spectroscopy circumvents the need for physical isolation or workup of the mixture components in the two samples. Using quantitative 1D ^1^H NMR, it is validated that the HSQC_0_ approach effectively removes all the undermining effects that plague quantification in common 2D NMR experiments, including differential attenuation due to relaxation in the two phases, pulse imperfections, partial decoupling, off-resonance effects, and incomplete coherence transfer in the case of scalar coupling variation. These results should be of widespread interest as partitioning into biomolecular condensates is crucial for the calibration of computational physicochemical models of phase separation and key to the further understanding of cellular biochemistry involving membraneless organelles.

## Introduction

Liquid-liquid phase separation (LLPS) is a fundamental organizing principle of the cell, facilitating the compartmentalization of biomolecules in the absence of membrane-bound structures.^1,2^ These compartments, often referred to as biomolecular condensates, harness the principles of LLPS to create specialized environments, where favorable self-interactions between biological polymers outweigh the entropic impetus towards mixing.^3–5^ Condensates are often highly dynamic and show properties that are normally associated with liquid droplets such as fusing, wetting, and dripping. Condensates also modulate enzymatic reaction rates, as they create distinct chemical environments that are sequestered from their surroundings, and yet permeable enough to allow the diffusion of molecules inside and between them. This has led to the suggestion that condensates can act as reaction crucibles for biochemical reactions inside the cell.^6,7^ Once a condensate forms, a partitioning equilibrium is established with molecules in the surroundings like that between water and an immiscible solvent. The partitioning coefficients of proteins in the two phases vary widely, with concentration imbalances that can extend up to 1,000-fold.^8–10^

The study of small molecule partitioning between the biomolecular condensates and their surrounding aqueous phase, referred to as dense and dilute phase, respectively, provides valuable insights into the microenvironment within these condensates. Furthermore, it is a crucial step in determining the biochemical activities of a condensate. However, partitioning coefficients for small molecules have been measured for only a handful of systems.^11–13^ Currently, most studies measure partitioning coefficients by comparing the fluorescence intensity from confocal fluorescence microscopy, where the dense and dilute phase can be determined simultaneously.^3,5,11,14,15^ This microscopy modality has the advantage of being able to measure partitioning of molecules into individual condensates inside cells but requires that they are either intrinsically fluorescent or can be labelled with a fluorophore. The latter is not feasible for many small molecule metabolites and drugs without adversely affecting their partitioning. An alternative approach using chromatography-coupled mass spectrometry was therefore developed and the partitioning coefficients for ∼1700 small molecules were quantified.^11^ Regrettably, only qualitative agreement was demonstrated for a subset accessible to fluorescence microscopy, and large differences (some as large as 100-fold) were observed for the partitioning of many compounds. This suggests that neither method is a “gold standard” and highlight a need for better analytical methods for measuring partitioning of small molecules into condensates. Access to comprehensive certified experimental data is also key for validating and optimizing molecular mechanics forcefields as well as training machine learning methods for predicting partitioning across condensates with different properties.

As an alternative, we therefore explore here an orthogonal approach based on quantitative nuclear magnetic resonance (qNMR) spectroscopy – a validated method for absolute and relative quantification.^16–19^ For example, in one-dimensional (1D) qNMR spectroscopy the intrinsic quantitative nature stems from the signal integral being proportional to the number of contributing nuclei in the sample, as long as considerations are made to starting from equilibrium (Zeeman) polarization. This requires that the inter-scan relaxation delay is sufficiently long to allow for full recovery by spin-lattice relaxation, with or without the assistance of a paramagnetic co-solute.^20,21^

Unfortunately, in many instances, one-dimensional NMR spectroscopy is not sufficient. Such is the case of the phase-separating systems studies here, where the background signals of the host proteins that constitute the condensates as well as carbohydrates, that can act as crowders^22^, strongly interfere with such an analysis, as illustrated in Fig. 1.

**Figure 1.**
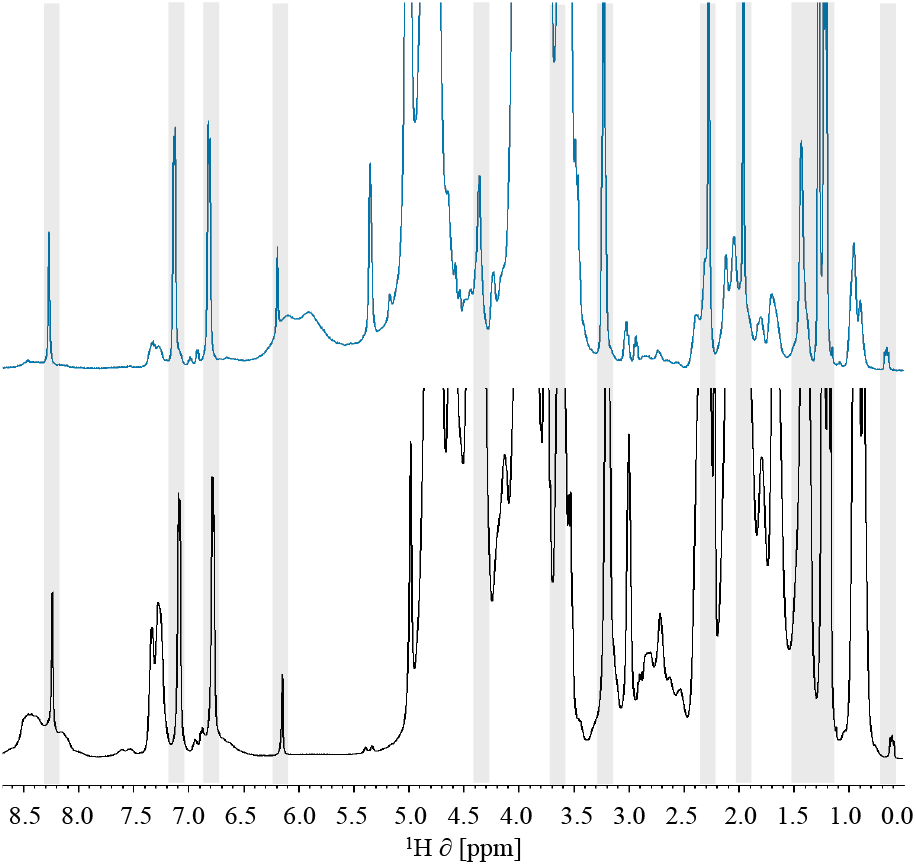
1D proton NMR spectra of six small molecules dissolved in the dilute (blue) and dense (black) parts of a phase-separated system made from fish gelatin and dextran. The NMR signals stemming from the six dissolved compounds are highlighted in gray.

In this situation, 2D NMR experiments can overcome this congestion hurdle and provide the necessary resolution. However, how to deal with the signal attenuation in the two (very unalike) phases during an entire 2D pulse sequence is not trivial. Over the years, various methods have been proposed that focus on pulse sequence performance improvements, analytical approaches to correct for magnetization evolution and relaxation, or a combination of these.^23–25^ One approach to improve quantification of the 2D HSQC pulse sequence was made by Heikkinen and coworkers, and coined “quantitative HSQC” (Q-HSQC).^26^ Herein, the strategy was to remove the effects of J_CH_ couplings by averaging four HSQC experiments with different delays during the INEPT period. The Q-OCCA-HSQC pulse sequence was later proposed to introduce further improve quantification: CPMG pulse trains in the INEPT and rINEPT periods to mitigate J_HH_ coupling evolution, composite CPMG pulse trains to reduce off-resonance effect on ^13^C, and adiabatic pulses during t_1_-evolution.^27,28^ More recently, an alternative approach was proposed by Castañar and coworkers, called perfect-HSQC, containing “perfect echo” INEPT modules during transfer periods to suppress J_HH_.^29^ However, none of these approaches avoid or quantify relaxation losses - particularly during magnetization transfer periods – a unique issue when investigating small molecules in condensates, as the relaxation times can change significantly due to differences in the physical properties of the phases. It is therefore necessary to account for the relaxation differences in the different phases.

A solution to the above stalemate can be forged by using the ^1^H-^13^C “extrapolated time-zero HSQC” (Heteronuclear Single Quantum Coherence spectroscopy), referred to as HSQC_0_, an experiment that has previously been used for concentration determination of metabolites in dilute solution.^30–32^ Its strength lies in the ability to remove the action of the entire pulse sequence through extrapolation, such that relaxation, imperfect pulses and decoupling, off-resonance effects, and coherence transfer mismatch are all removed simultaneously and effectively. In brief, the method works by recording spectra where the entire pulse sequence has been concatenated between one to *i* times. The signal amplitude A of peak *n* in the 2D ^1^H-^13^C HSQC_0_ can be expressed using Equation (1),

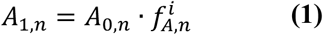

The amplitude attenuation factor, 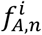 represents the signal attenuation due to *i* concatenations of the entire 2D pulse sequence prior to acquisition. Spectra recorded with *i* = 1,2,3, … will cause the signal to be attenuated by the same factor each time it is repeated. Hence, by extrapolation to *i* = 0, the integrated signal amplitude, *A*_0,*n*_, is obtained - *as if the pulse sequence was never applied*. The attenuation factor is specific for peak *n* and is obtained by extrapolation of the individual signal integrals. The integrated intensity after *i* repetitions can be recast in linear form, Equation (2):

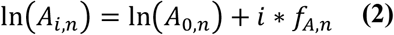

which makes it easy to obtain the integrated signal from a simple linear regression. The absolute concentration of the molecules in a sample is then obtained by dividing the integrated intensity, *A*_0,*n*_, with the number of protons in the peak *n* and comparing it to an internal reference with known concentration. It should be kept in mind, however, that due to relaxation, it may only be possible to repeat the pulse sequence a limited number of times *i*.

Using the integrated signal amplitudes derived from the extrapolation as relative concentrations, the partition coefficients (PCs) are then computed using the following definition:

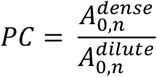

## Results and discussion

### Validation of the ^1^H^-13^C HSQC_0_ method for quantification of compounds in a mixture

We first validated the HSQC_0_ approach using a collection of six compounds in D_2_O. The panel consisted of compounds with different physicochemical properties and display a small number of idiosyncratic NMR signals to preclude signal overlap (see Table I).

**Table I.** Compounds dissolved in D_2_O to test quantification. The compounds, chemical shifts, and results from the extrapolation to HSQC_0_ are given.

| Compound | Signal | $\delta$ $^1\text{H}$<br>[ppm] | Extrapolated $\ln(A)$<br>[a.u.] | $R^2$ |
| --- | --- | --- | --- | --- |
| p-cresol | ring CH | 7.09 | $19.58 \pm 0.05$ | 0.9996 |
| p-cresol | ring CH | 6.77 | $19.51 \pm 0.08$ | 0.9987 |
| p-cresol | CH<sub>3</sub> | 2.22 | $20.01 \pm 0.01$ | 0.9999 |
| cAMP | sugar CH | 6.15 | $18.51 \pm 0.15$ | 0.9938 |
| 1,4-dioxane | CH<sub>2</sub> | 3.75 | $21.14 \pm 0.01$ | 0.9999 |
| ethyl-guanidine sulfate | CH<sub>2</sub> | 3.18 | $20.01 \pm 0.04$ | 0.9996 |
| ethyl-guanidine sulfate | CH<sub>3</sub> | 1.17 | $20.46 \pm 0.02$ | 0.9999 |
| *tert*-butanol | CH<sub>3</sub> | 1.24 | $21.28 \pm 0.04$ | 0.9997 |
| DSS | CH<sub>3</sub> | 0.00 | $19.12 \pm 0.17$ | 0.9960 |

As one-dimensional NMR spectroscopy is a gold standard for quantitative analysis^16,19,33^, we used a fully relaxed 1D ^1^H NMR spectrum as reference (see Supporting Information, Fig. S1). This requires an interscan delay of 5*T_1_, and therefore the longest ^1^H T_1_ was determined by inversion recovery.^20^ We found for this sample that the longest T_1_ was 1.08 s for 1,4-dioxane (example curves and T_1_ times are provided in the Supporting Information, Fig. S2 and Table S1). In what follows, an interscan delay of 7 s was therefore used, and reference signal intensities for all peaks were determined by peak integration.

Subsequently, the HSQC_0_ method was applied to the same sample and extrapolated intensities were determined by linear fitting (see Supporting Information, Table S2). Extrapolations for all compounds showed errors below 2.5%. A test for self-consistency was additionally made by comparing different signals from the same compound, and this showed uncertainties of maximum 4%. The quantifiability of the HSQC_0_ method was then compared to signal intensities from the quantitative 1D experiment (single excitation pulse) described above and proved that the HSQC_0_ method is highly quantitative (ρ = 0.996), as illustrated in Fig. 2.

**Figure 2.**
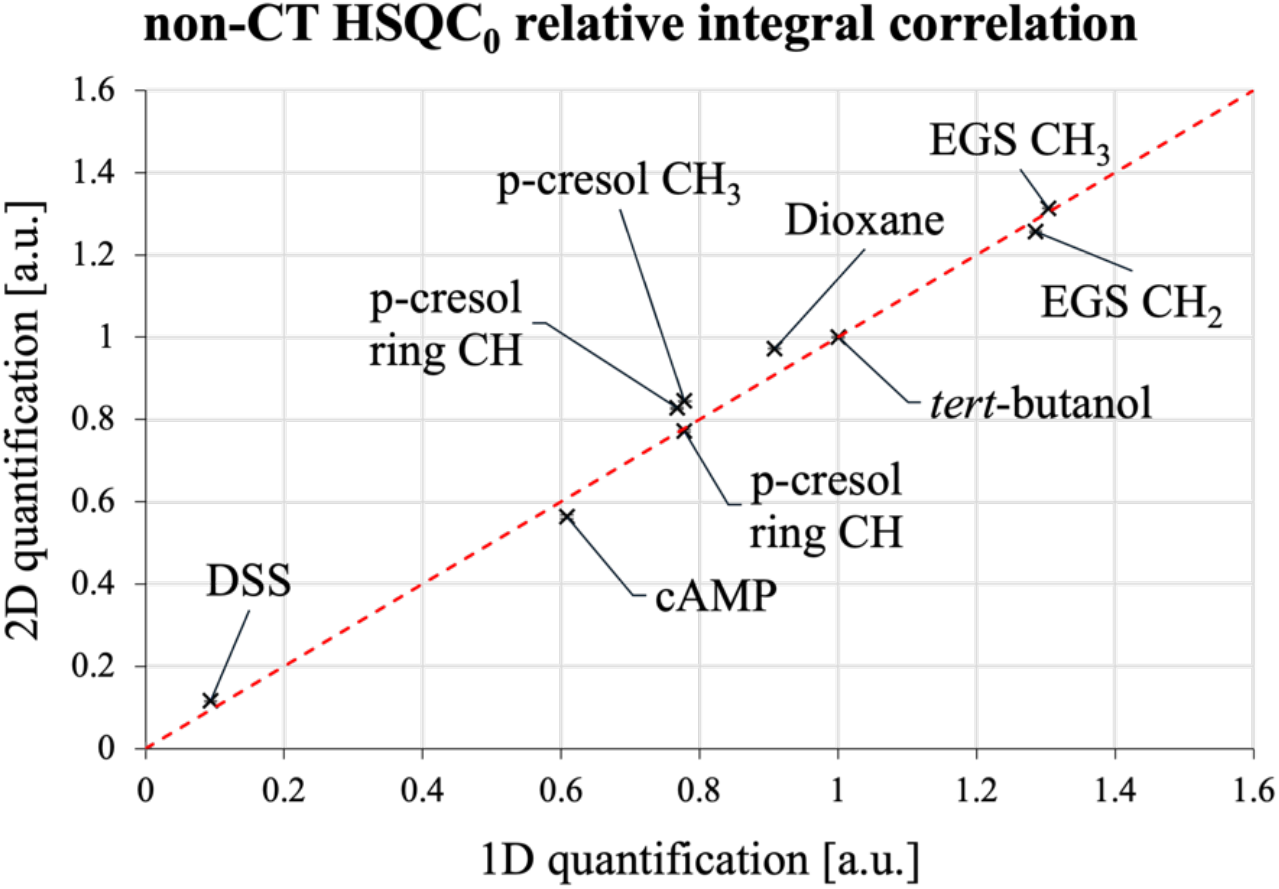
Compound quantification by HSQC_0_ compared to 1D ^1^H qNMR. Signals are normalized with respect to tert-butanol.

### Partitioning coefficients determined in a liquid-liquid phase separated system

Next, to mimic the physical conditions of a protein-based liquid-liquid phase-separated (LLPS) system, we mixed dextran and cold-water fish gelatin, a model condensate described by Riedstra & McGorty.^34^ As the host scaffolds are made of proteins and carbohydrates at high concentration, it presents a real-life challenge for NMR spectroscopy, as the client signals occur in the same regions. The small molecules (Table I) were added to the dextran-gelatin mixture, and after mixing the system was left for 24 hours to phase separate. The phases were separated upon preparation of the NMR samples. The NMR spectra of dilute and dense phase are shown in Figure 1. Signal overlap renders 1D quantification impractical, prompting us to resort to the extrapolated HSQC_0_ method. To ensure proper quantitation for the two-phase system, T_1_ relaxation times were measured for the small molecules in both phases (see Supporting Information, Table S1). Overall, relaxation is faster in the dense protein-rich phase than in the dilute sugar-rich phase, and faster than in pure D_2_O, as expected due to increased viscosity.

The HSQC_0_ experiments result in well-resolved spectra for both phases (see Supporting Information, Fig. S3). Examples of signal extrapolation for three signals are shown in Figure 3. As anticipated, a clear loss in intensity is observed when using more repetitions of the pulse sequence, primarily due to relaxation and uncertainty in the integrals grows as clearly evinced for the HSQC_3_ experiment. While the error increase is notable for all signals, the fitting still provides good linear correlations and fits with low errors. Using the extrapolated integrals from dense and dilute phases, partition coefficients were determined for eight signals and are listed in Table II. Two signals, dioxane and p-cresol, had overlapping signals with the dense phase in the 2D HSQC spectrum and could therefore not be interpreted accurately. Since some compounds give rise to multiple signals, it is possible to check for internal consistency of the determined partitioning coefficients. Two estimates were available for p-cresol, ethyl-guanidine, and cAMP, with the highest variation being 6%. All partition coefficients were larger than 1, meaning that they preferentially partitioned to the dense phase, except for *tert*-butanol, which showed no preference.

**Table II.** Extrapolated HSQC_0_ values for small molecules in the dilute and dense phases of a dextran-gelatin model system and derived partitioning coefficients.

| Signal | Extrapolated ln(A)<br>Dilute phase [a.u.] | Extrapolated ln(A)<br>Dense phase [a.u.] | Partition coefficient<br>[dense/dilute] |
| --- | --- | --- | --- |
| p-cresol ring | 19.54 ±0.02 | 19.91 ±0.05 | 1.45 ±0.07 |
| p-cresol CH<sub>3</sub> | 19.86 ±0.02 | 20.19 ±0.01 | 1.39 ±0.01 |
| cAMP | 17.67 ±0.07 | 18.07 ±0.08 | 1.49 ±0.09 |
| cAMP CH | 17.31 ±0.20 | 17.66 ±0.13 | 1.42 ±0.10 |
| dioxane<sup>1</sup> | -- | -- | -- |
| ethyl-guanidine<br>sulfate CH<sub>2</sub> | 19.86 ±0.03 | 20.06 ±0.02 | 1.22 ±0.04 |
| ethyl-guanidine<br>sulfate CH<sub>3</sub> | 20.24 ±0.04 | 20.38 ±0.04 | 1.15 ±0.01 |
| *tert*-butanol | 20.74 ±0.03 | 20.73 ±0.02 | 0.99 ±0.01 |
| DSS | 18.89 ±0.20 | 19.13 ±0.02 | 1.27 ±0.09 |

<sup>1</sup> The signal for dioxane was not sufficiently resolved for reliable integrals to be obtained.

**Figure 3.**
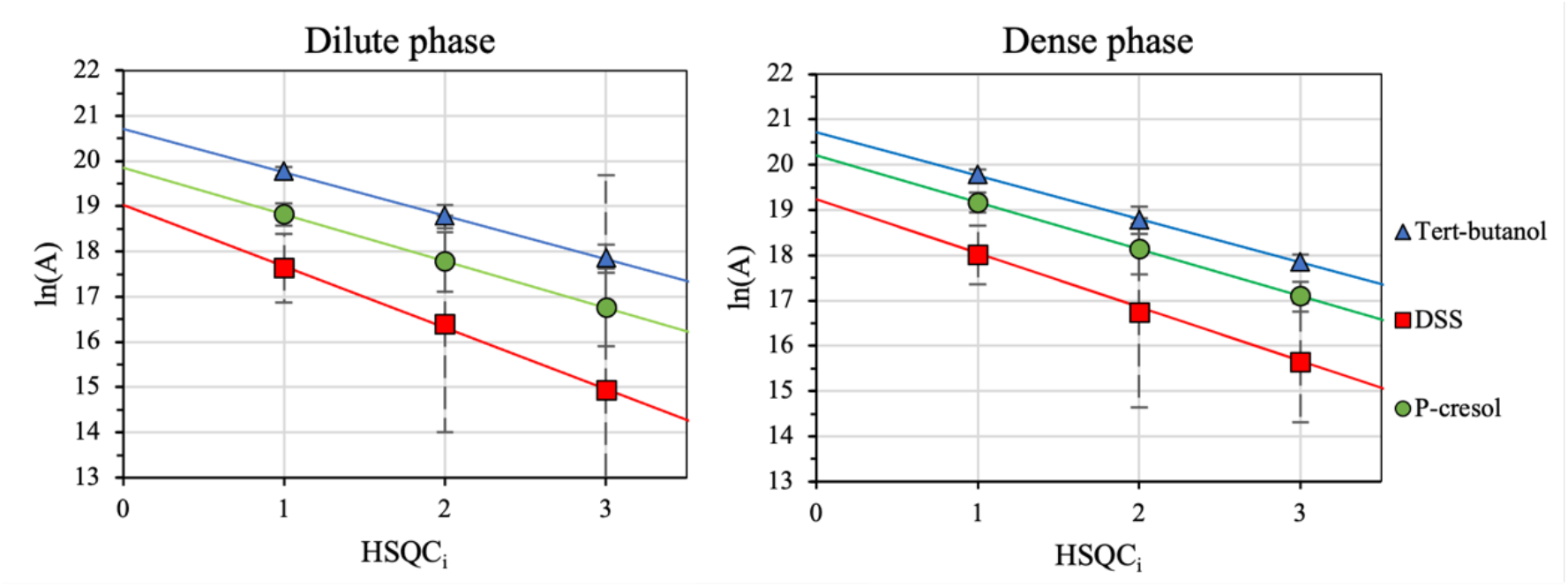
Quantification in the dilute and dense phases of three compounds in the dextran-gelatin model system by HSQC_0_.

It is demonstrated that the ^1^H-^13^C HSQC_0_ method is well-suited for quantification of metabolites in the two phases formed by a gelatin/dextran mixture. Since most cellular metabolites contain at least one carbon-bound hydrogen atom, this method promises to be a general method, as it solves the resolution limitation of ^1^H 1D qNMR, where overlap with the phase-generating molecules effectively precludes its general application. For four test compounds, DSS, p-cresol, cAMP, and ethyl-guanidine, similar partitioning coefficients ∼1.2-1.4 were obtained, with the concentration of each molecule being slightly higher in the dense (protein-rich) phase.

The method proved quantitative for the more viscous environment of the dense phase. While the sensitivity in the third repetition, HSQC_3_, is low, partitioning coefficients are determined with less than 7% error, using the multiple signals of individual compounds as an internal control. Although some condensates are known to have apparent viscosities for macromolecules that are more than 10 times that of water, the presented method may still be applicable for small molecules that do not interact too strongly with the macromolecular matrix.^35^ We anticipate that this method may be widely applicable for studying small-molecule partitioning in phase-separating systems that suffer from background interference of the matrix background.

## Conclusion

In this study, we set out to investigate the applicability of 2D qNMR for the determination of partitioning coefficients in biomolecular condensates. We successfully measured the division of a panel of small molecules in the two phases of a model system that have physiochemical properties of liquid-liquid phase separation of biomolecular condensates. Since molecules typically give rise to several NMR signals, it was also possible to test for internal consistency. Quantification turned out to be very high and was typically within 2%, indicating that this method may serve as a reliable approach for small molecule partitioning across a wider variety of artificial and biological condensates to understand their behavior and function.

## Acknowledgements

We thank Dr. Milo Westler, Dr. Kaifeng Hu, and Prof. Dr. John Markley (University of Wisconsin) for sharing Bruker NMR pulse sequences. The work presented in this article is supported by Novo Nordisk Foundation grant NNF20OC0063808, ‘BOUNDLESS’ and the Villum Foundation (grant 27963). Access to the NMR spectrometers at the Danish Center for Ultrahigh-Field NMR Spectroscopy (Ministry of Higher Education and Science grant AU- 2010- 612-181) and expert help from Dr. Dennis Wilkens Juhl are gratefully acknowledged.

## Experimental section

### Small molecule NMR samples

A mixture of small molecules with different physiochemical properties was produced. The mixture contained the following compounds in D_2_O: tert-butanol, p-cresol, 3,5-cyclic adenosine monophosphate, 1,4-dioxane, ethyl-guanidine sulfate and DSS (4,4-dimethyl-4-silapentane-1-sulfonate). The solution was buffered with 20 mM Na-phosphate pH 7.4 and 0.5 mM gadoteridol was included to enhance metabolite spin relaxation.^21^

### Dextran-gelatin NMR samples

Aqueous solutions of 500 kDa dextran 19 % by weight and cold-water fish gelatin 12 % by weight in 20 mM NaH_2_PO_4_ at pH 7.4 buffer were prepared. The ratio of dextran and gelatin was 3:2 to ensure enough material of both phases. To the mixture were added the following compounds: tert-butanol, p-cresol, cyclic AMP, 1,4-dioxane and ethyl- guanidine sulfate, to final concentrations near 20 mM. DSS (4,4-dimethyl-4-silapentane-1- sulfonate) was included as chemical shift reference standard at a ten-fold lower amount. The mixture was centrifuged at 7000 rpm for 5 minutes and left at 21°C for 20 hours. The two phases were separated, and 0.5 mM gadoteridol (to enhance relaxation rates) and 10% v/v D_2_O (for the NMR lock) were included in the final samples for NMR spectroscopic analysis.

### NMR spectroscopy

All NMR experiments were recorded at 298K at 500 MHz ^1^H frequency using a room temperature probe head, and processed using TopSpin v. 3.5.7.^36^ A quantitative 1D ^1^H spectrum was acquired with an interscan delay (d1) of 60 s. 128204 complex datapoints with a spectral width of 16 ppm and 1024 scans per FID were used. Line broadening of 0.30 Hz was applied to 1D data. Phase correction was performed manually, and the baseline was autocorrected. Manual peak integration was done with TopSpin (Bruker Biospin). Error estimation of the integrals was done using the standard deviation obtained for 10 empty baseline regions.

### Proton T_1_ determination NMR experiments

Inversion recovery experiments to determine T_1_ of the small molecules in D_2_O and the model system were performed using *t1ir* and an adapted version of *noesygppr1d* pulse sequence for inversion recovery^20^ respectively. The following settings were used for the *t1ir* experiments: 32768 complex data points with a spectral width of 12 ppm, 32 scans and d1 = 30s. T_1_ relaxation was determined using 16 randomly ordered delays ranging from 0.0001 to 30 seconds. The following settings were used for the *noesygppr1d*- based inversion recovery experiment: 16024 complex data points with a spectral width of 16 ppm, 8 scans and d1 = 1s. Additionally, trim pulses of 1ms were applied prior to the interscan delay (d1) to ensure equivalent saturation at the end of each FID. T_1_ relaxation was determined using 10 randomly ordered delays ranging from 0.01 to 12 seconds. Curve fitting was done with the T1T2 module in Topspin.^36^

### Time-zero HSQC experiments

Four versions provided by the research group of Prof. J. Markley were tested for the extrapolated time-zero HSQC method.^30^ The first two used P- and N-type selection with either constant-time evolution or regular t_1_ evolution, whereas the other two used States-TPPI acquisition. The HSQC_0_ experiments recorded using P- and N-type selection and regular (*i*.*e*. non-constant time) t_1_ evolution is chosen for quantification experiments. Three experiments with increasing repetition of the HSQC block, i.e., HSQC_1_, HSQC_2_ and HSQC_3_, were performed. The experiments for the dense and dilute phase samples were measured consecutively, on samples that exceed the coil height to ensure that an identical volume of spins is excited. The carrier frequencies were set to 4.7 (^1^H) and 75.0 ppm (^13^C), respectively. 2048 × 256 complex data points with spectral widths of 19 ppm and 170 ppm with 16 scans per FID and an interscan delay of 7s was used. Data were processed to 8192 × 1024 datapoints with a shifted-sine-bell function applied in both domains. Baseline and phase correction were done manually. DSS was used as chemical shift reference (direct reference).

## References

1. Alberti, S. & Dormann, D. Liquid–Liquid Phase Separation in Disease. Annu. Rev. Genet. 53, 171–194 (2019).

2. Hyman, A. A., Weber, C. A. & Jülicher, F. Liquid-Liquid Phase Separation in Biology. Annu. Rev. Cell Dev. Biol. 30, 39–58 (2014).

3. Dumelie, J. G. et al. Biomolecular condensates create phospholipid-enriched microenvironments. Nat. Chem. Biol. 20, 302–313 (2024).

4. Alberti, S., Gladfelter, A. & Mittag, T. Considerations and Challenges in Studying Liquid-Liquid Phase Separation and Biomolecular Condensates. Cell 176, 419–434 (2019).

5. Kilgore, H. R. & Young, R. A. Learning the Chemical Grammar of Biomolecular Condensates. Nat. Chem. Biol. 18, 1298–1306 (2022).

6. Zhang, Y., Narlikar, G. J. & Kutateladze, T. G. Enzymatic Reactions inside Biological Condensates. J. Mol. Biol. 433, 166624 (2021).

7. O’Flynn, B. G. & Mittag, T. The role of liquid–liquid phase separation in regulating enzyme activity. Curr. Opin. Cell Biol. 69, 70–79 (2021).

8. Diamond, A. D. & Hsu, J. T. Protein partitioning in PEG/dextran aqueous two-phase systems. AIChE J. 36, 1017–1024 (1990).

9. Huddleston, J., Abelaira, J. C., Wang, R. & Lyddiatt, A. Protein partition between the different phases comprising poly(ethylene glycol)-salt aqueous two-phase systems, hydrophobic interaction chromatography and precipitation: a generic description in terms of salting-out effects. J. Chromatogr. B. Biomed. Sci. App. 680, 31–41 (1996).

10. Kato, S., Garenne, D., Noireaux, V. & Maeda, Y. T. Phase Separation and Protein Partitioning in Compartmentalized Cell-Free Expression Reactions. Biomacromolecules 22, 3149–3624 (2021).

11. Ambadi Thody, S. et al. Small-molecule properties define partitioning into biomolecular condensates. Nat. Chem. 16, 1794–1802 (2024).

12. Bannan, C. C., Calabró, G., Kyu, D. Y. & Mobley, D. L. Calculating Partition Coefficients of Small Molecules in Octanol/Water and Cyclohexane/Water. J. Chem. Theory Comput. 12, 4015–4024 (2016).

13. Sepehri, B., Drew, K. & Villegas, J. A. Come for the atmosphere, stay for the interactions: Deciphering small molecule partitioning into biomolecular condensates. Cell Chem. Biol. 30, 1337–1339 (2023).

14. Kilgore, H. R. et al. Distinct chemical environments in biomolecular condensates. Nat. Chem. Biol. 20, 291–301 (2024).

15. Abbas, M., Lipiński, W. P., Wang, J. & Spruijt, E. Peptide-based coacervates as biomimetic protocells. Chem. Soc. Rev. 50, 3690–3705 (2021).

16. Bernstein, M. et al. qNMR: The Handbook. 2nd edition, BoD - Books on Demand (2024).

17. Giraudeau, P. Quantitative 2D liquid-state NMR. Magn. Reson. Chem. MRC 52, 259–272 (2014).

18. Martineau, E.Dumez, J.-N. & Giraudeau, P. Fast quantitative 2D NMR for metabolomics and lipidomics: A tutorial. Magn. Reson. Chem. MRC 58, 390–403 (2020).

19. Giancaspro, G. et al. The qNMR Summit 5.0: Proceedings and Status of qNMR Technology. Anal. Chem. 93, 12162–12169 (2021).

20. Mulder, F. A. A., Tenori, L., Licari, C. & Luchinat, C. Practical considerations for rapid and quantitative NMR-based metabolomics. J. Magn. Reson. 352, 107462 (2023).

21. Mulder, F. A. A., Tenori, L. & Luchinat, C. Fast and Quantitative NMR Metabolite Analysis Afforded by a Paramagnetic Co-Solute. Angew. Chem. Int. Ed Engl. 58, 15283–15286 (2019).

22. Xie, S., Yue, C., Ye, S. & Li, Z. Biological Condensate Growth: Examining the Impact of Solute Crowder on Size Expansion. Biomacromolecules 26, 323–331 (2025).

23. Lane, D. et al. Assessing the potential of quantitative 2D HSQC NMR in 13C enriched living organisms. J. Biomol. NMR 73, 31–42 (2019).

24. Rai, R. K., Tripathi, P. & Sinha, N. Quantification of Metabolites from Two-Dimensional Nuclear Magnetic Resonance Spectroscopy: Application to Human Urine Samples. Anal. Chem. 81, 10232–10238 (2009).

25. Bingol, K., Zhang, F., Bruschweiler-Li, L. & Brüschweiler, R. Quantitative Analysis of Metabolic Mixtures by Two-Dimensional 13C Constant-Time TOCSY NMR Spectroscopy. Anal. Chem. 85, 6414–6420 (2013).

26. Heikkinen, S., Toikka, M. M., Karhunen, P. T. & Kilpeläinen, I. A. Quantitative 2D HSQC (Q-HSQC) via Suppression of J-Dependence of Polarization Transfer in NMR Spectroscopy: Application to Wood Lignin. J. Am. Chem. Soc. 125, 4362–4367 (2003).

27. Koskela, H. Use of NMR techniques for toxic organophosphorus compound profiling. J. Chromatogr. B 878, 1365–1381 (2010).

28. Koskela, H., Kilpeläinen, I. & Heikkinen, S. Some aspects of quantitative 2D NMR. J. Magn. Reson. 174, 237–244 (2005).

29. Castañar, L., Sistaré, E., Virgili, A., Williamson, R. T. & Parella, T. Suppression of phase and amplitude JHH modulations in HSQC experiments. Magn. Reson. Chem. MRC 53, 115–119 (2015).

30. Hu, K., Westler, W. M. & Markley, J. L. Simultaneous quantification and identification of individual chemicals in metabolite mixtures by two-dimensional extrapolated time-zero 1H-13C HSQC (HSQC0). J. Am. Chem. Soc. 133, 1662–1665 (2011).

31. Hu, K., Ellinger, J. J., Chylla, R. A. & Markley, J. L. Measurement of absolute concentrations of individual compounds in metabolite mixtures by gradient-selective time-zero 1H-13C HSQC with two concentration references and fast maximum likelihood reconstruction analysis. Anal. Chem. 83, 9352–9360 (2011).

32. Hu, K., Wyche, T. P., Bugni, T. S. & Markley, J. L. Selective Quantification by 2D HSQC0 Spectroscopy of Thiocoraline in an Extract from a Sponge-Derived Verrucosispora sp. J. Nat. Prod. 74, 2295–2298 (2011).

33. Bharti, S. K. & Roy, R. Quantitative 1H NMR spectroscopy. TrAC Trends Anal. Chem. 35, 5– 26 (2012).

34. Riedstra, C. P. & McGorty, R. Liquid–Liquid Phase Separation: Undergraduate Labs on a New Paradigm for Intracellular Organization. The Biophysicist 1, (2019).

35. Wei, M.-T. et al. Phase behaviour of disordered proteins underlying low density and high permeability of liquid organelles. Nat. Chem. 9, 1118–1125 (2017).

