## Supplementary material for "Quantification of small molecule partitioning in a biomolecular condensate with 2D NMR spectroscopy": SI

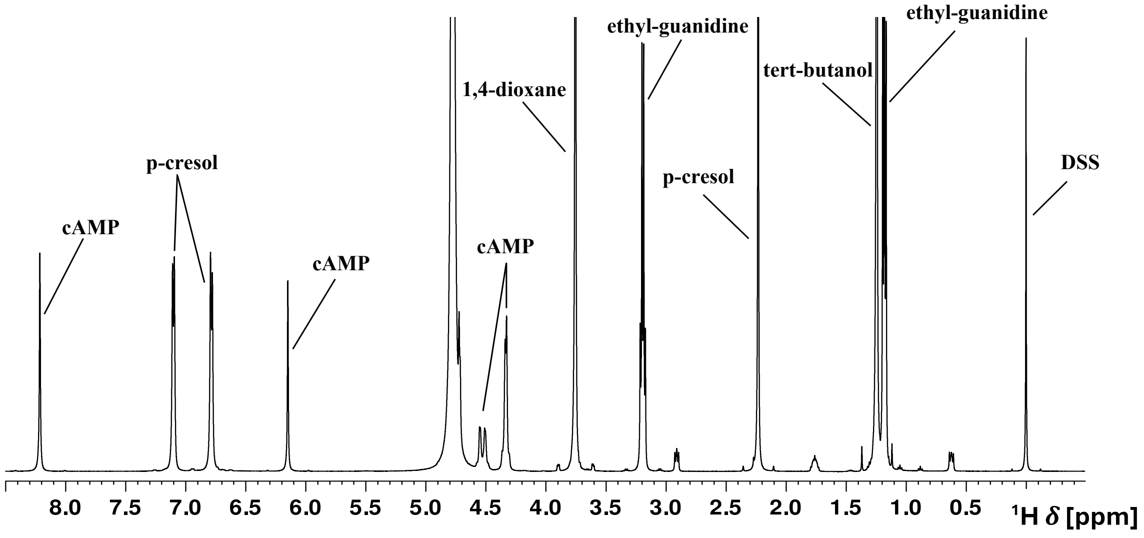


**Figure S1.** $^1$H 1D spectrum in $D_2O$ used for quantification. Signals are indicated with corresponding compound.

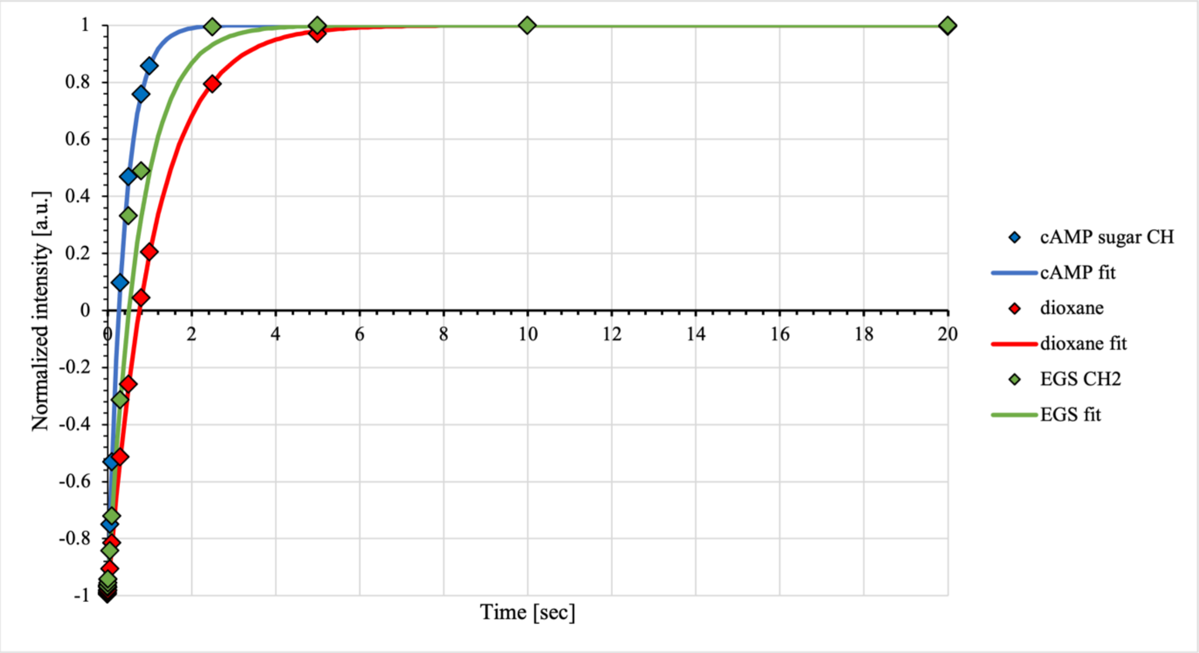


**Figure S2.** Inversion recovery data and fitted $T_1$ recovery curves for selected signals.

**Table S1.** $T_1$ times for all samples.

| Compound | $^1H$ ∂ [ppm] | $D_2O$ [sec] | Dense phase [sec] | Dilute phase [sec] |
|---|---|---|---|---|
| p-cresol ring CH | 7.09 | -- | 0.209 ±0.003 | 0.271 ±0.001 |
| p-cresol ring CH | 6.77 | -- | 0.183 ±0.004 | 0.235 ±0.003 |
| cAMP | 6.15 | 0.377 ±0.004 | 0.259 ±0.007 | 0.357 ±0.005 |
| 1,4-dioxane | 3.75 | 1.080 ±0.0025 | 0.552 ±0.008 | 0.689 ±0.003 |
| Ethylguanidine $CH_2$ | 3.18 | 0.735 ±0.002 | 0.415 ±0.004 | 0.375 ±0.006 |
| p-cresol | 2.22 | -- | 0.262 ±0.001 | 0.310 ±0.002 |
| *tert*-butanol | 1.24 | 0.883 ±0.005 | 0.591 ±0.002 | 0.632 ±0.002 |
| Ethylguanidine $CH_3$ | 1.17 | 0.831 ±0.003 | 0.513 ±0.002 | 0.543 ±0.004 |
| DSS | 0.00 | 0.795 ±0.004 | 0.536 ±0.003 | 0.629 ±0.002 |

**Table S2.** Extrapolated intensities from $HSQC_0$ approach for small molecule mixture in $D_2O$.

| Compound | $^1H$ ∂ [ppm] | Extrapolated ln(A) [a.u.] | $R^2$ |
|---|---|---|---|
| p-cresol ring CH | 7.09 | 19.58 ±0.05 | 0.9996 |
| p-cresol ring CH | 6.77 | 19.52 ±0.08 | 0.9987 |
| cAMP | 6.15 | 18.5 ± 0.2 | 0.9938 |
| 1,4-dioxane | 3.75 | 21.14 ± 0.01 | 0.9999 |
| Ethylguanidine $CH_2$ | 3.18 | 20.01 ±0.04 | 0.9996 |
| p-cresol | 2.22 | 20.01 ± 0.01 | 0.9999 |
| *tert*-butanol | 1.24 | 21.28 ± 0.04 | 0.9997 |
| Ethylguanidine $CH_3$ | 1.17 | 20.455 ± 0.02 | 0.9999 |
| DSS | 0.00 | 19.1 ± 0.2 | 0.9960 |

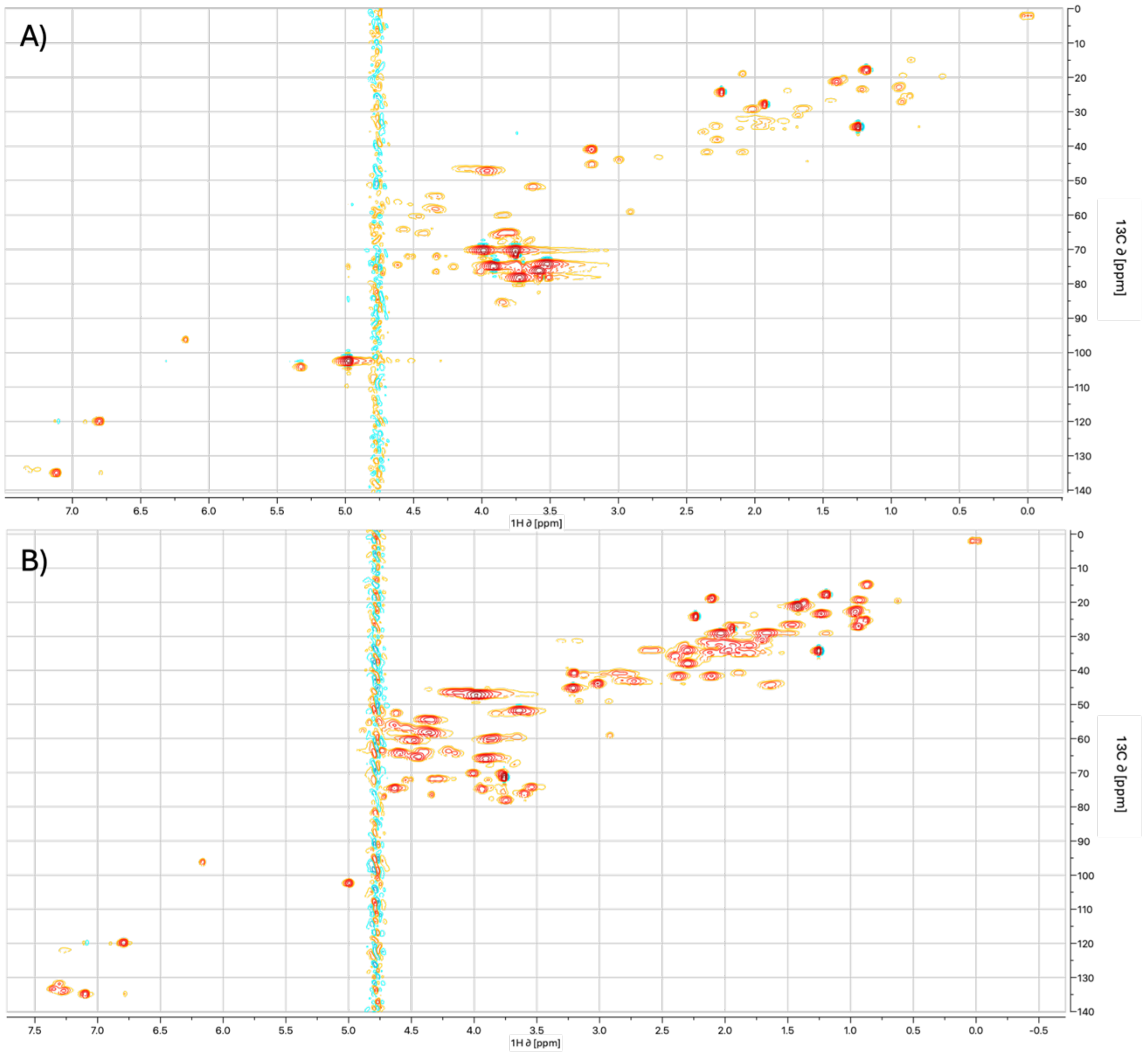


**Figure S2.** $HSQC_0$ spectra of A) dilute and B) dense phase of the model condensate.
